# Enabling stable coexistence by modifying the environment

**DOI:** 10.1101/676890

**Authors:** Alejandro Pastor, Juan Carlos Nuño, José Olarrea, Javier de Vicente

**Affiliations:** Department of Applied Mathematics in Aerospace Engineering. Universidad Politécnica de Madrid – ETSIAE, Madrid, Spain; Department of Applied Mathematics. Universidad Politécnica de Madrid, Madrid, Spain

## Abstract

In this work coexistence is studied using a model based on two classical population models: the quasispecies of Eigen [1] and the daisyworld presented by Watson and Lovelock [2]. It is assumed that species are able to modify the environment. We show that this ability enables the coexistence between, at most, two species in equilibrium. Given an initial population, the problem arises as to determine which of the many equilibrium populations, i.e. extinction, only one species or coexistence of two species, will be reached as a function of the species characteristics, specifically their capacity to modify the environment and the optimal growth rate. These results are obtained under general assumptions, which broadens its applicability to other fields as evolutionary biology and social sciences.

## 1 Introduction

The question of whether an initial population formed by different species will persist on time is of great relevance in Ecology and Evolutionary Biology [3–5]. However, to get conclusive results seems to be elusive because of the so many factors that are involved in the dynamic behavior of the population [6]. Besides, the individual properties of the species have to be confronted with initial and boundary conditions.

From a dynamical point of view, this question is translated to which asymptotic equilibrium the population will attain when multiple stable equilibria exist. Classical qualitative analysis, using local properties of equilibria, is not enough to solve this question [7, 8]. Finding its solution requires a global approach that considers the properties of all the species as well as the possible interactions among them and the influence of the environment. Unfortunately, despite the large number of papers devoted to study the relationship between biodiversity and stability in ecosystems [3, 5], there are no standard techniques to analyse this kind of complex systems [9].

Our model is founded on two classical models: the Quasispecies model introduced by Eigen in the earlies seventies [1] and the so called Daisyworld model presented by Watson and Lovelock in 1983 to study the homeostatic properties of ecosystems [2, 10]. From the first, we use the dynamic description, the concept of species fitness and the population constraint that brings about a selective process. From the second, the hypothesis that the individuals have the ability to influence their environment. The model we present here belongs to a different category of those where the environment changes independently of the population (see, for instance, [11, 12])

There are several examples where the species ability to modify their environment are specially relevant. Tumor cells are able to modify the surrounding tissues by segregating chemical substances that increase the fitness of malign cells [13]. Soil modification by microbial communities is being reported as one of the main driving factors of these ecosystems [14]. Finally, climate change caused anthropogenically, i.e. induced by human actuations, constitutes a dramatic example of the effect of modifying the temperature by the species that inhabit the planet [15].

In this paper we study a population of replicators that are able to modify their environment [16]. The environment is here described by an unique scalar variable, that we will call temperature. We assume that species fitness depend on this temperature that changes over time as a function of the population distribution. Maximum fitness for each species is reached at a particular optimal temperature value, decreasing when temperature departs from this value. Species influence on temperature can be either positive or negative and it is not coupled with their optimal temperature. Besides, total population is bounded by a carrying capacity of the system. As a consequence, the initial population undergoes a selective proccess that ends in an equilibrium population formed by the survival species, if any.

It is worth noting that, as a consecuence of the feedback between the population and its environment, the fitness of each species varies with time. There is no proper way to rank the species out of the particular context they are placed.

For the sake of clarifying, the main assumptions of the model are listed below:

i. A carrying capacity exists that bounds the total population,
ii. The environment is described by a unique scalar function, its temperature, that is a function of time,
iii. The influence of each species on the temperature is proportional to its population size,
iv. The fitness of each species is a symmetrical function of temperature with a single peak at its optimal temperature,
v. Species only interact with each other indirectly through this temperature and the resource (space) constraint.

The next section presents in detail the mathematical model. Results concerning with the stability analysis of population of low diversity are obtained in the third section. Fourth section presents the results obtained from the simulations of populations with large initial biodiversity. We conclude and discuss these results, as well as their implications, in the final section.

## 2 The mathematical model

We consider a population of error-free self-replicative species (replicators) *I_i_* for *i* = 1,…,*S*. The size of the total population at time *t* is *N*(*t*) and, can be determined as the sum of the population of each phenotype *N_i_*(*t*):

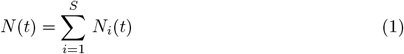

Species *I_i_* has associated a fitness function *f_i_* that can be described by two real numbers: *T_i_* and *α_i_*. The first one, *T_i_*, stands for its optimal growth temperature whereas, *α_i_* denotes its influence on the environment. Both parameters, *T_i_* and *α_i_*, give a measure of the survival probability of *I_i_* in each generation. The relative fitness *f_i_* depends on the rest of species of the population through the global temperature *T*. In particular, we assume that the relative fitness of copy *I_i_* is given by

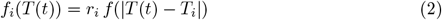

where *r_i_* is a non-negative parameter and *f*(*x*) is a function that exhibits an unique maximum at *x* = 0 and decreases monotonically to 0 as *x* → ∞ (see Figure 1).

**Fig 1.**
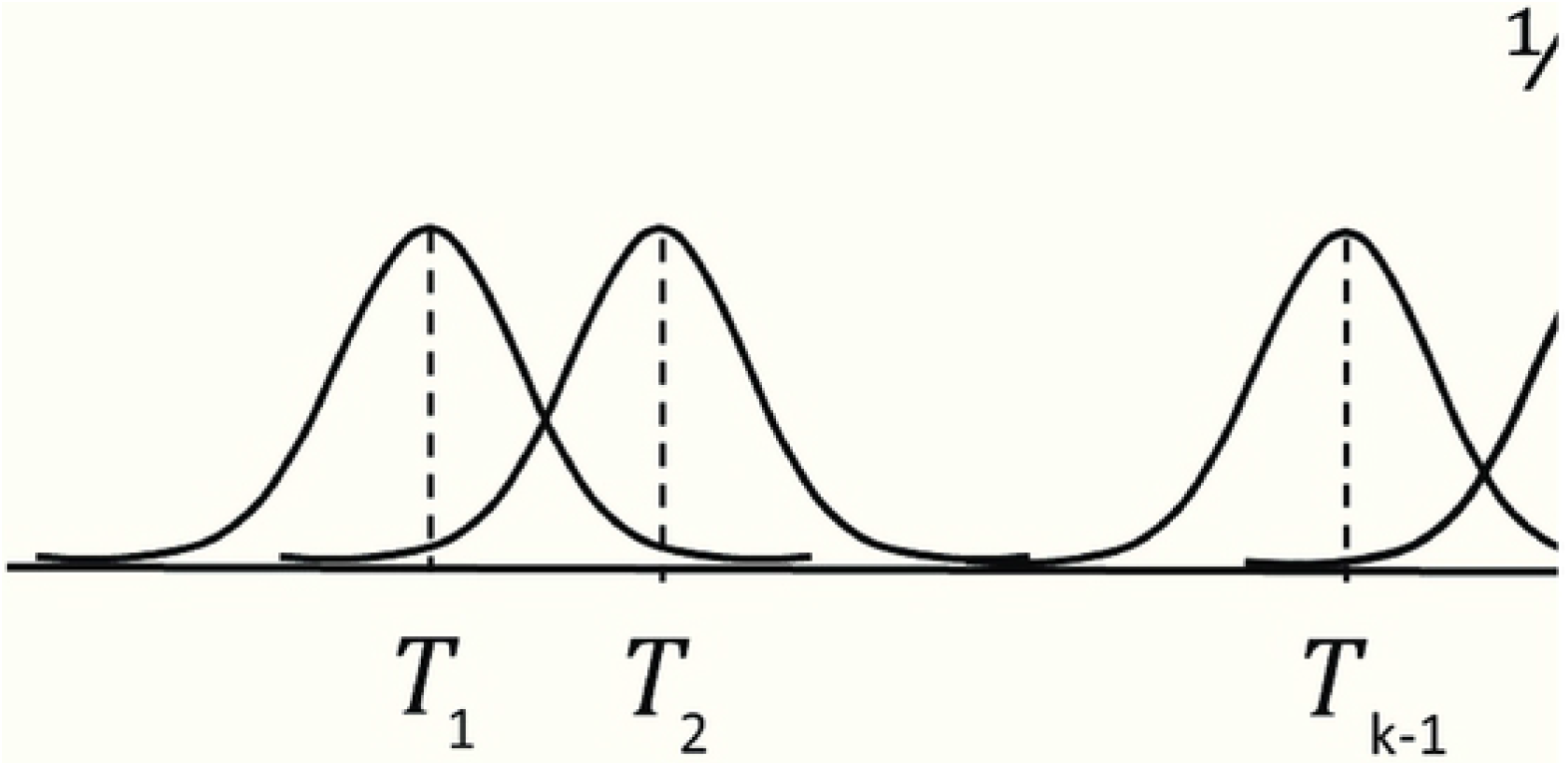
Examples of possible *f*(*x*) in fitness equation (2).

The intensive parameter *T* characterizes the environment. We suppose that each species *I_i_* has a linear influence on *T*, weighted by the real parameter *α_i_*. Assuming that external perturbations that could modify the value of *T* are negligible, the time evolution of the global temperature is given by:

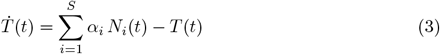

In order to induce a selective process, we assume that the system has a maximum carrying capacity *K*, the upper bound of the total population. Let the function of time, *s*(*t*), be the available space at time *t*:

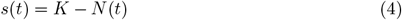

then, the global fitness of species *I_i_* is given by:

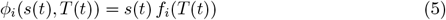

At each time step, the reproduction rate of every species is a function of both the population size and the external temperature and becomes null as the population size approaches its maximum capacity *K*.

The time evolution of each species population *N_i_* can be described by a system of Ordinary Differential Equation (ODE):

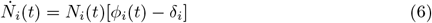

for *i* = 1, 2,…,*S*. The parameter *δ_i_* is the death rate of species *I_i_*. For sake of simplicity, if not explicitely indicated, we will assume in what follows that *r_i_* = *r_j_* ≡ *r* and *δ_i_* = *δ_j_* ≡ *δ* for all *i, j*. Under these assumptions, species differ each other by their optimal temperature *T_i_* and their capacity to modify the external temperature *α_i_*. Notice that, as all species have the same *r* value, we can redefine the fitness function *f* (see 2) including the common factor *r* in it, i.e. from now on:

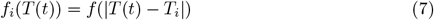

It is convenient to normalize the equations with respect to the carrying capacity, *K*. Let 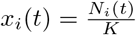 for *i* = 1,…,*S*. In so doing, we have to redefine the system parameters adequately: *r*\* = *r K* and 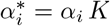, although, we keep the same notation in the equations:

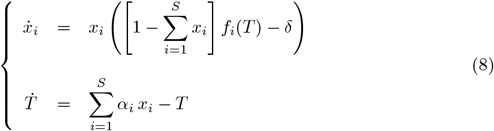

The non-linear character of this ODE system prevents its analytical solution. Nevertheless, a complete qualitative analysis for different cases has been carried out and it is presented in the following.

First of all let us find the equilibrium points. There are two posibilities to cancel the differential equation for species *x_i_* in system 8. Either *x_i_* = 0, or the parenthesis on the right vanishes, i.e.,

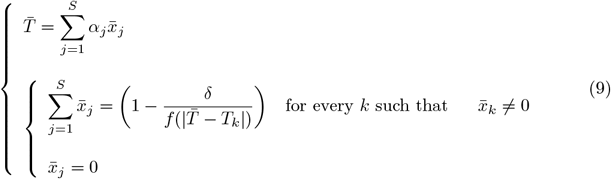

The local stability character of each of these equilibrium points is given by the Jacobian matrix associated to system 8, evaluated on them.

For every 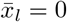, the l-row in the Jacobian matrix has only one non-null entry at the diagonal position *J_l, l_*. Consequently, the associated corresponding eigenvalues are:

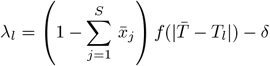

In order to compute the rest of the eigenvalues, we only consider the ODEs that correspond to the values 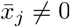. If we arrange these variables in the *m* last equations, we form a non-trivial box of dimension (*m* + 1) × (*m* + 1) in the lower right part of the Jacobian. The equilibrium points of this subsytem verify:

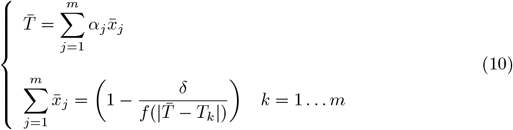

Note that this system of algebraic equations is inconsistent for large values of *m*. In practice, coexistence of more than two species is impossible, except in degenerate cases of species with the same optimal temperature, which we will consider indistinguishable. It is straightforward to prove that if *T_r_* = *T_s_* for a pair of species we can redefine the system using linear combinations of *x_r_* and *x_s_* in such a way that one of the combinations is stationary and the other behaves as a new species with an *α* value that is a combination of *α_r_* and *α_s_* and the system behaves exactly as the one we are studying with one variable less.

When a solution exists, the associated Jacobian is straightforwardly computed. The entry *J_k, j_* para *k, j* = 1 … *m* is given by:

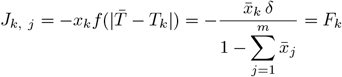

The entries of the last column are (∀*k* = 1,…,*m*):

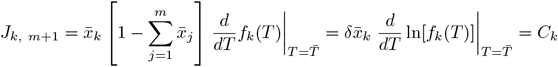

and the last row:

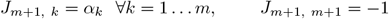

With this notation, the structure of the Jacobian is:

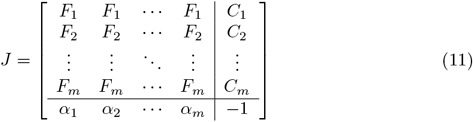

In the following sections we will study separately each of the equilibria and determine their stability conditions over the parameters, particularly, on *T_i_* and *α_i_*.

## 3 Fixed environment: *α_i_* = 0 for all *i*

The reference case, that reduces the ODE system (8) to the classical error-free quasispecies model, corresponds to the situation when the species have no capacity to modify the temperature, i.e. *α_i_* = 0 for all *i*. According to equation (8) temperature decreases exponentially from its initial condition to zero. When the stationary value has been reached, the fitness of all the species remains constant in time: *f_i_* = *f*(|*T_i_*|) and our model replicates the quasispecies model. It can be proven that asymptotic coexistence is not possible and only two non-degenerate equilibria exist: (i) Extinction, i.e. all the species of the population die out and (ii) Selection of only one species, whereas the rest disappear. The equilibrium of *m* different species requires the *m* species to comprise the same fitness, i.e. *f_i_* = *f_j_*, which is a degenerate situation. Obviously, neutral situations in which more than one species have the same fitness are possible but these species are considered as indistinguishable in this paper.

The largest fitness corresponds to the species with optimal temperature *T_i_* closer to zero, *f*(*min*|*T_i_*|) and, therefore this species (lets assume index *k*) is the only one that can take over the whole population. Its equilibrium size is given by:

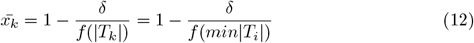

From the qualitative analysis explained in previous section one can inferred that, as *α_i_* = 0 for all *i*, the last row of the Jacobian has only one non-null entry, *J*_*S*+1,*S*+1_ = −1. Besides, for 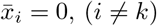 the corresponding rows in the jacobian matrix have again only a non-null entry, *J_i,i_*. So the eigenvalues directly emerge:

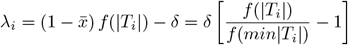

As *f* is a monotonically decreasing function, all these eigenvalues are negative. The row of the Jacobian corresponding to each surviving species reads as follows:

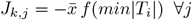

and the corresponding eigenvalue is then:

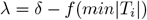

If *f*(*min*|*T_i_*|) < *δ* all eigenvalues are negative ant this is the asymptotically stable state, whereas if *δ* < *f*(*min*|*T_i_*|), the final system tends towards extintion.

## 4 Promoting coexistence by modifying the environment

Let us consider the case when the species are able to modify the environment (*α_i_* ≠ 0) and analyze the different possible stationary states. As we stated before, contrary to the quasispecies model, now coexistence equilibria of at most two species can occur.

### 4.1 Extinction

We consider first the equilibrium point: 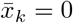 for all *k* = 1,…, *S* and 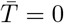, i.e. the extiction of the whole population. The analysis is similar to the preceding case *α_i_* = 0 ∀*i* and it is again straightforward to prove that this equilibrium point is asymptotically stable if

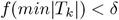

### 4.2 Only one survival

Let us assume that only one species has a population different from 0, for instance *I_k_*. That is, 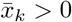 and 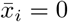 for all *i* ≠ *k*. The equilibrium point is obtained from the system:

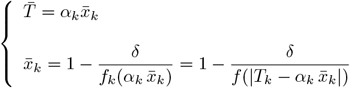

As 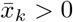

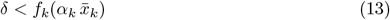

The eigenvalues associated to *i* ≠ *k* are

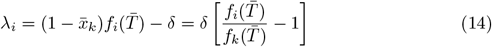

The condition for this equilibrium point to be asymptotically stable is:

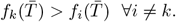

This last condition implies that:

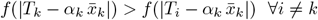

which means that the value of 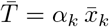 must be closer to *T_k_* than any other *T_i_*.

At this point, as we will see later, it is convenient to reorganize the species according to their optimal temperatures, satisfying

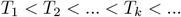

in a increasing succession as depicted in Figure 1.

The former condition reads now:

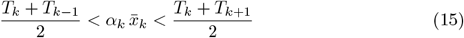

The rest of eigenvalues are those of the submatrix in the Jacobian:

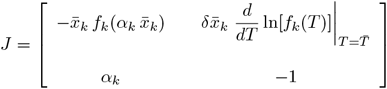

The associated characterisitic polynomial is:

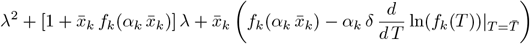

The equilibrium point is asymptotically stable if the two roots have negative real parts. This occurs when:

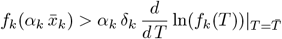

### 4.3 Coexistence

Let us assume that the unique two survival species are *I_k_* and *I_m_*. The values of the equilibrium population are the solution of the algebraic system:

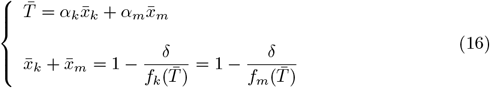

The first obvious condition that must be satisfied is 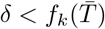.

The right part of the second equation is satisfied if 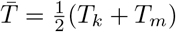 or *T_k_* = *T_m_*. The second case can be reduced to the case studied in the previous section, again by using linear combinations of these two species. One of the new variables is stationary and the other behaves as a new species with an *α* value that is a linear combination of *α_k_* and *α_m_*.

Let us focus on the first one: 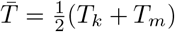. Eigenvalues associate to *i* ≠ *k, m* are:

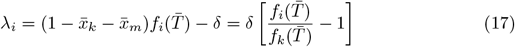

Again, the condition for this equilibrium point to be asymptotically stable is:

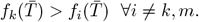

Then,

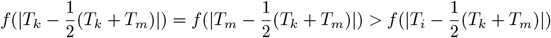

which means that the value of 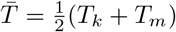 must be closer to *T_k_* and *T_m_* than any other *T_i_* and, therefore, *T_m_* y *T_k_* must be consecutives, i.e. *m* = *k* + 1 (Hence the convenience of arranging the species according to their optimal temperatures, as we stated before).

Now, the resulting linear system:

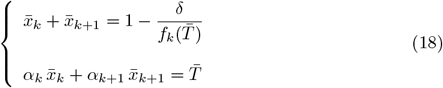

can be straightforwardly solved yielding:

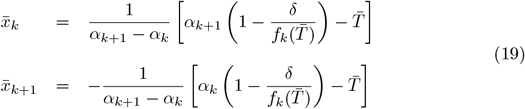

For these two concentrations to be simultaneously positive 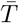 must have a value in between 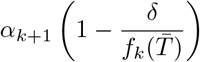 and 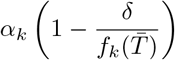.

As it will be proven next, asymptotical stability is discarded in the case *α*_*k*+1_ > *α_k_*. For *α*_*k*_ > *α*_*k*+1_, as both values in equation 19 must be positive 19, equation 18 implies:

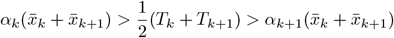

The remaining three eigenvalues associated to this coexistence equilibrium are obtained from the reduced Jacobian matrix:

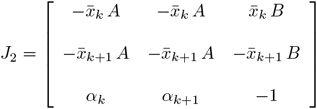

where

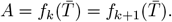

and

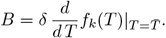

Notice that *A* > 0 and *B* < 0 as *f_k_* is a decreasing function at 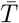 (see Figure 1). The characteristic polynomial associated to *J*_2_ is:

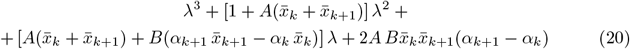

The Routh-Hurwitz criterion requires the following conditions to assure three roots with negative real parts:

1. All coefficients are positives. This implies that: *α_k_* > *α*_*k*+1_ as stated before. In addition,

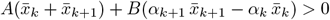
2. 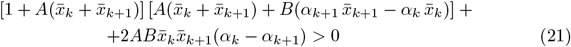

These general conditions can be applied to seek examples where species coexistence exist. In particular, as shown in the next section, we will find examples where multi-stability between equilibria of coexistence appear and, consequently, the final population depends on the initial conditions.

### 4.4 Four species model

In order to illustrate the results obtained in this section, we explore the numerical solution of several four species cases where multi-stability appears. This model of four initial species exhibits, among others, two kind of multi-stabilities that do not occur in the classical models: (i) bi-stability between two different coexistence equilibrium points and (ii) tri-stability among two single species equilibria and one coexistence point.

The first situation occurs for the species described in Table 1. The rest of parameters are *r* = 1 and *δ* = 0.1. The fitness function chosen for all the cases from now on reads

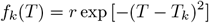

**Table 1.** 4-species model 1

| $\alpha$ | $T_{op}$ |
| --- | --- |
| -0.6 | -0.9 |
| -1 | -0.8 |
| 0.1 | -0.1 |
| -0.2 | 0.2 |

Among other equilibria, this model has the following two asymptotically stable equilibrium populations: (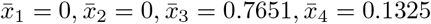, *T* = 0.05) and (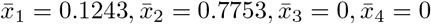, *T* = −0.85). The final equilibrium these four species will achieve depends on the initial population. In Figure 2 we show the projection on the plane Π (whose coordinate axis are *x*_1_ + *x*_2_ and *x*_3_ + *x*_4_) of the solution obtained for 1500 initial conditions, all with *T*(0) = 0. As it can be seen, the basin of attraction of each 2-coexistence is diffusively separated by the hyperplane *x*_1_ + *x*_2_ = *x*_3_ + *x*_4_ that is projected as a line in the plane Π.

**Fig 2.**
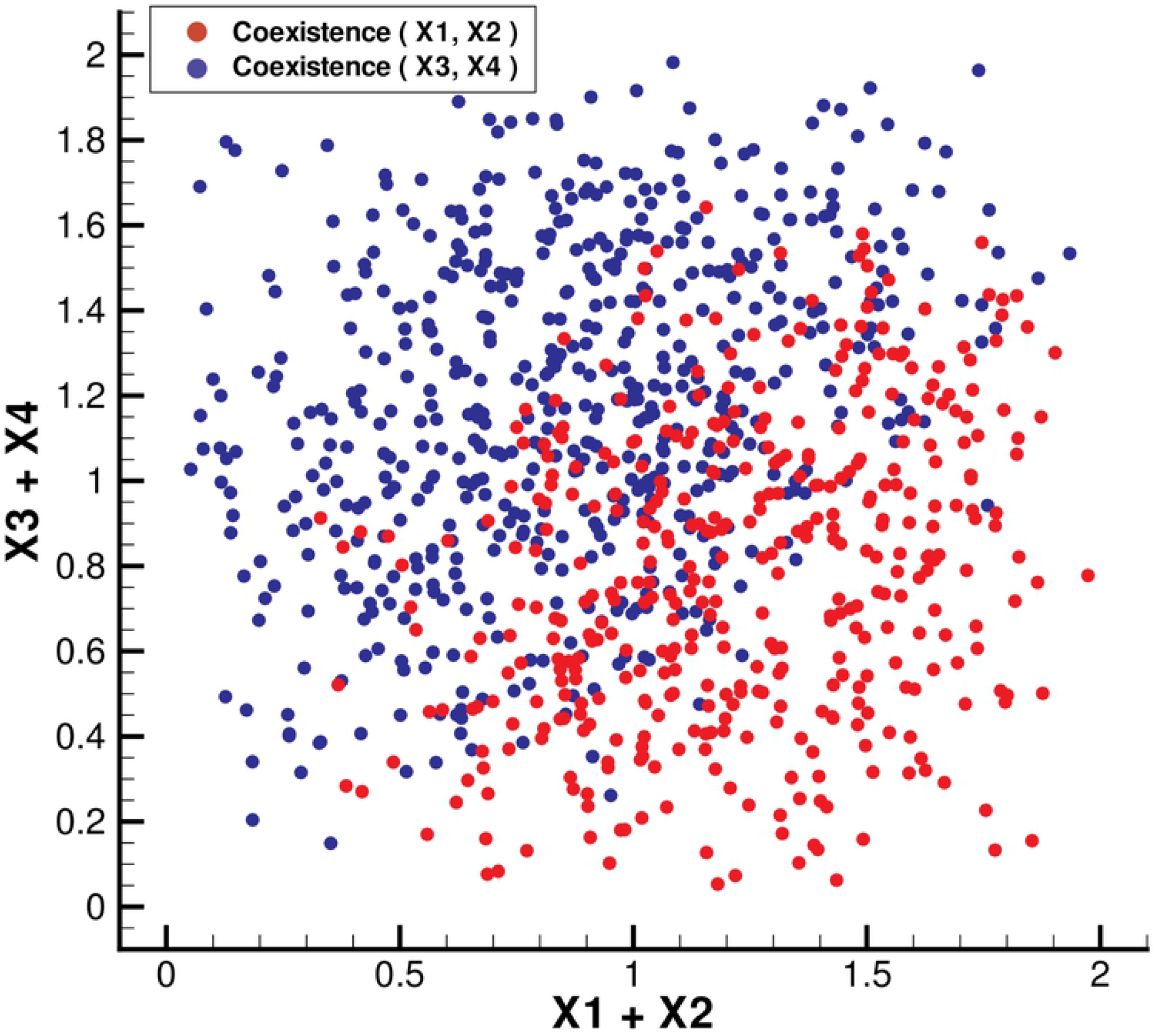
In order to describe the basins of attraction of the two 2-coexistence equilibrium solutions in the four species model with the parameter setup given in Table 1, we show a set of 1500 initial conditions and mark with red circles those that converge to the 2-coexistence between species *I*_3_ and *I*_4_ and with blue circles those initial conditions that converge to the 2-coexistence of species *I*_1_ and *I*_2_. The rest of the parameter values are *r* = 1 and *δ* = 0.1.

Another example of a case with four species that exhibits tri-stability is depicted in Table 2. The rest of the parameters is fixed as before.

**Table 2.** 4-species model 2

| $\alpha$ | $T_{op}$ |
| --- | --- |
| -1 | -0.9 |
| -0.7 | -0.85 |
| 0.1 | -0.1 |
| -0.2 | 0.2 |

Note that this example is a slight variation of the previous one, sharing the last two species. Nevertheless, among others equilibria, this model exhibits an unusual tri-stability among the following equilibrium points (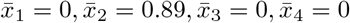, *T* = −0626), (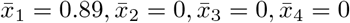, *T* = −0.9) and (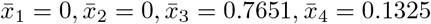, *T* = 0.05). This latter equilibrium already exists in the previous example.

In both cases, depending on the initial conditions the population tends to one of these equilibrium points. A tridimensional projection of 1000 initial conditions is depicted in Figure 3 where the coordinates axis are: *x*_1_, *x*_2_ and *x*_3_ + *x*_4_. As it can be observed, the coexistence equilibrum has the largest basin of attraction (blue points). On the other extreme, the single species equilibrium with 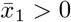 has the smallest basin of attraction (only seven red circles corresponding to 1-existence 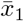 ocurrences out of 1000 initial conditions).

**Fig 3.**
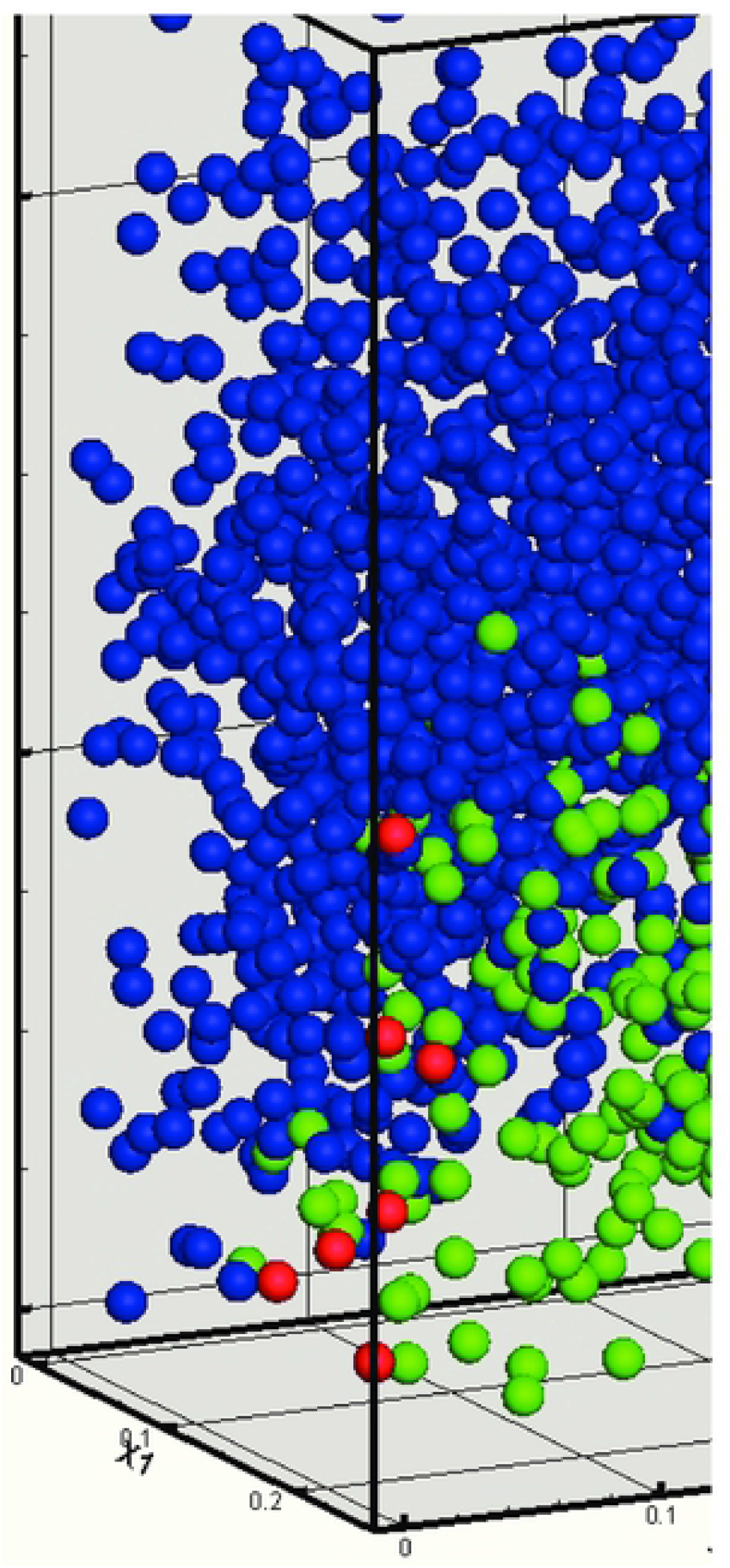
As in the previous figure, we show the basins of attraction of the three equilibria that exist in the four species model presented in Table 2 a 2-coexistence and two 1-coexistence. A set of 1500 initial conditions are classified depending on their final equilibrium population: blue circles correspond to the 2-coexistence formed by species *I*_3_ and *I*_4_, red circles to the 1-coexistence of species *I*_1_ and green circles to the 1-coexistence of species *I*_2_. The other parameter values are: *r* = 1 and *δ* = 0.1.

## 5 Populations with a large number of species

In the previous section, we have studied the stability properties of a system formed by few species, concretely, four. We have proven the existence of conditions that yield multistability between single species and coexistence equilibria. Under these conditions, the asymptotic equilibrium population depends on the initial populations of each of the species that form the population. In this section, we explore further this dependence in populations formed by a larger number of species. An important result to be pointed out is that coexistence equilibria with more than two species does not exist. We want to show in this section how the asymptotic behavior of this kind of populations depends on three factors: (i) the number of species, (ii) the initial conditions and (iii) the confluence of the properties of the species that initially form the population, i.e. their optimal temperature *T_k_* and the rate of influence, *α_k_*, over the environment temperature *T*.

### 5.1 Dependence with the number of species

In order to study the influence of the number of species that initially form the population, we carry out a numerical integration of the system of differential equations that describes the time evolution of each of the species population, *x_k_*(*t*) (see 8). As before, we assume that each species *I_k_* is characterized by *T_k_* and *α_k_*. The values of these parameters are taken randomly from intervals whose length is changed in each case. The initial populations of all the species are equal and their sum reaches half of the total population, i.e. 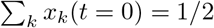. The rest of the parameters that define the system are taken as before, i.e. *r* = 1, *δ* = 0.1. For each number of species and for each *α*-interval and *T*-interval, we perform 30 numerical integrations and quantify the equilibrium populations. The optimal temperature and the rate of influence of the initial species are randomly chosen from two given intervals that are symmetric with respect to 0: *T_i_* ∈ [−*T_max_, T_max_*] and *α_i_* ∈ [−*α_max_, α_max_*].

Figure 4 depicts the curves of the proportion of coexistence equilibria for the case *α*-interval = [−1, 1] for different *T*-intervals. The rest of the proportion corresponds to single species equilibria except for the large *T*-intervals and lower number of species where the proportion of extinction is significant (see Figure 4). For the other *T*-intervals this proportion is null. As it can be seen, the occurrence of 2-coexistence equilibria is more probable for population formed initially by a large number of species. This tendency is more effective when the *T*-interval is shorter. On the contrary, this proportion is reduced to half when the *T*-interval is [−10, 10] even for large number of species.

**Fig 4.**
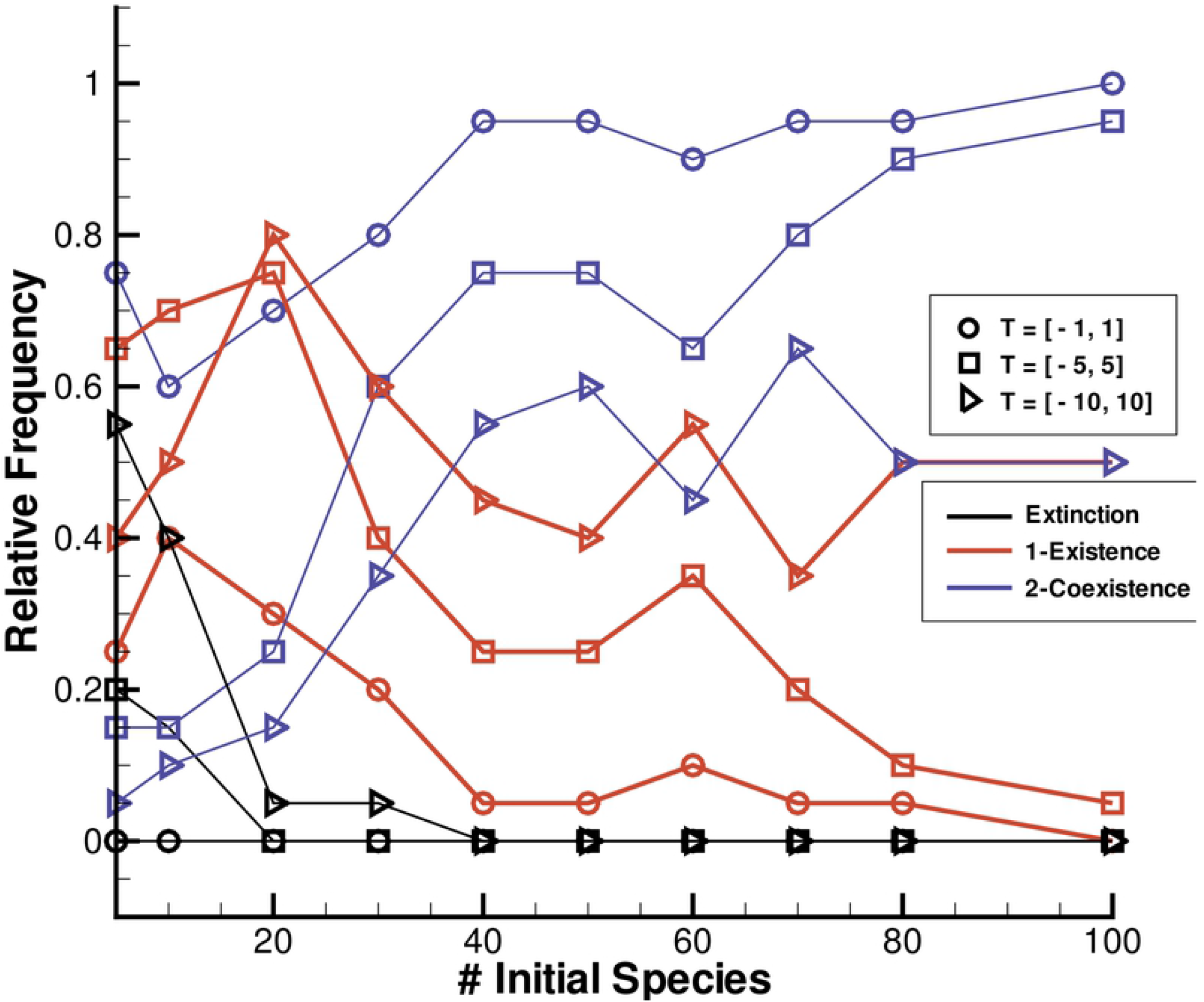
Relative frequency of the coexistence equilibria as a function of the number of initial species for different *T*-intervals. The interval of variation of *α_k_* is [−1, 1]. An initial population of *N* species with random values of *T_k_* and *α_k_* in these intervals is considered. All the species are equally represented in the initial population that occupies half of the carrying capacity. The initial external temperature is *T*(0) = 0. Each point in the curves is the average over 30 realizations.

Similar results are obtained when the *T*-interval is fixed to [−10, 10] and we vary the length of *α*-interval as it is summarized in Table 3. In that table the probability of reaching a 2-coexistence equilibrium as a function of the number of initial species is shown. In this case, the probability of extinction is null and, consequently, the probability of survival of only one species for each number of initial species is 1 minus the value given by the corresponding curve. As in the previous figure, it can be seen that the coexistence equilibria are more likely to occur in populations with larger number of species. This effect is more important when the *α*-interval is larger.

**Table 3.** Frequency of the 2-coexistence equilibrium point for different intervals of variation [−*α, α*] as a function of the number of species *S*. The interval of variation of the optimal temperatures of the species is [−10, 10]. An initial population is selected of *S* species with random values of *T_k_* and *α_k_* in these intervals. All the species are equally represented and the sum of their sizes occupies half of the initial population and the initial external temperature is *T*(0) = 0. Each value is the average over 30 realizations.

| α-interval | Number of initial species | | | | | | | | | |
| --- | --- | --- | --- | --- | --- | --- | --- | --- | --- | --- |
|  |  | 5 | 10 | 20 | 30 | 40 | 50 | 60 | 80 | 100 |
| $[-0.1, 0.1]$ | | 0.15 | 0.2 | 0.35 | 0.15 | 0.45 | 0.5 | 0.55 | 0.85 | 0.75 |
| $[-0.5, 0.5]$ | | 0.3 | 0.45 | 0.75 | 0.75 | 0.65 | 0.95 | 0.8 | 1 | 1 |
| $[-1, 1]$ | | 0.53 | 0.63 | 0.76 | 0.93 | 0.96 | 0.96 | 0.93 | 0.96 | 1 |
| $[-2, 2]$ | | 0.6 | 0.65 | 0.95 | 0.95 | 0.95 | 0.95 | 1 | 1 | 1 |
| $[-5, 5]$ | | 0.5 | 0.7 | 0.75 | 0.85 | 1 | 0.95 | 1 | 1 | 1 |
| $[-10, 10]$ | | 0.35 | 0.7 | 0.7 | 0.9 | 0.9 | 0.85 | 0.95 | 0.85 | 0.95 |

## 6 Discussion

The capability of some species to modify the environment has been postulated as a factor of stabilization that can promote biodiversity [3]. The coupling between biota and the environement could endowe the ecosystem with homeostatic properties that enables a better adaptation [5, 17]. The Gaia hypothesis is mainly based on this interaction [18, 19]. Indeed, Evolution favours those species that are able to keep the environment under specific conditions that are assumed good enough for Life.

A huge debate exists about the evolutionary occurrence of this kind of interactions between evolving species and an environment governed by the Laws of Physics. Hypothetically, biological species would be selected to fit particular environmental conditions. This would require, of course, a certain stability of the environment that, in evolutionary time, has not always occurred. However, our planet Earth has abruptly changed its physical properties from the origin of Life. Likely, most of the species that were selected in a static environment could be later displaced by new environmental conditions. Thus, it can be said that darwinian evolution is acting constrained by the dynamics of the environement. As a consequence, evolution selects species with the capability of controling the rapid and drastic changes of the environment in their own benefit. The problem is how to explain the fixation of a systemic property from the selective pressure on the individuals of the population [17, 20–22]. Even more challenging is to explain the existence of stable populations when mutation enables the appearance of species that can drive the environment towards destabilization.

This paper shows that when species are able to modify the environment, coexistence can be enhanced. As a matter of fact, for populations with a large number of species, coexistence is the most probable equilibrium population. However, if this modification is not coupled with the own species characteristics then, this capability does not assure its own persistence. The question about which species will survive under specific conditions can not be answered exclusively by carrying out a stability analysis. Qualitative stability analysis, for instance, based on the linearization around equilibria only provides local information about the dynamics of small perturbations: if the perturbation decays the equilibrium points are said to be asymptotically stable. On the contrary, if the perturbation increases over time the equilibrium point is said unstable. When the system exhibits multiple equilibrium points this analysis cannot solve the aymptotic behaviour when the population starts from a particular initial condition. Linear stability analysis says nothing about the attraction basin. Nevertheless, we have got a useful result, proven in section 4, that states a necessary condition that must satisfy two species that coexist in equilibrium: their temperatures must be consecutive. Unfortunately, this condition is not conclusive and further investigation is required to predict the species selected from a given initial population.

In order to solve this problem, we have applied a different approach that seeks to determine the equilibrium point (asymptotically stable) which is achieved from an initial population. In particular, we assume that the species optimal temperatures and their rate of influence on the environment are randomly taken. The population starts with a given number of species homogeneously populated and whose sum is well below the carrying capacity of the system *K*. After a transient period large enough to assure the system relaxation, we take note of the survival species and analyse their properties. We check that all these equilibria are asymptotically stable by a qualitative analysis. As expected, depending on the values of the external temperature the equilibrium population may differ. We note that most of the equilibrium populations are coexistence of two species that present a definite relationship between their optimal temperature *T_k_* and the rate of influence on the environment *α_k_*, concretely *T_k_ α_k_* < 0. Few simulations yield with the survival of only one species. None, as expected, with more than three species.

The deterministic description provided by Ordinaty Differential Equations (ODE) assumes a negligible internal noise, i.e. a large population size. This is the case when the carrying capacity *K* = 10^5^ and the initial population sizes are *x_i_*(0) = 5 × 10^2^ for *i* = 1,…,*S*, values that are applied in most of the computations. To check that this approach is correct we have also simulated the dynamic of the population by using an individual-based algorithm (data not shown). We simulate a discrete population in which each inidividual can be chosen either to replicate or to die according to its (global) fitness. We note that for this value of *K* the results of these simulations agree in all cases with the numerical integration of the ODE system. On the contrary, we have detected some differences for small values of *K*. This effect could be relevant when considering mutation because mutants populations are very small at the first stages after appearance.

## 7 Concluding remarks

The stability of populations is a key issue in Ecology. Grimm and Wissel (1997) stressed the large number of definitions that have appeared in Ecology [23]. These definitions are classified into six classes that resume the concepts frequently used in this field: constancy, resilience, persistence, resistance, elasticity, and domain of attraction [24]. In some point or another all of them appear in this work. Specifically, we handle with persistence when studying the species that remain in the population after a period of time. Mathematical stability, as shown in section 2, provides information about the local resilience and, in addition, enables to estimate the domain of attraction of equilibria [7]. In contrast, this stability analysis does not inform about the persistence of the species composition of an initial population that change over time due to the existence of multiple stable equilibria [25]. In other words, this approach can not predict the equilibrium point to which the system will tend. The consideration that species can modify the environment and, as a consequence, the composition of the population, includes an extra difficulty to perform a qualitative analysis and prevent an analytical solution of the problem. Numerical integration and simulations are instead appropiate tools for solving this problem as it is shown in this paper.

An important subject to be handled in a forthcoming paper concerns the evolutionary properties of this kind of ecosystems. We have assumed that the mutation rate of the species is null and consequently we have avoided the appearance of new species in the population. In this kind of models that consider the ability of species to modify the enviroment, the appearance of a new species with influence on the environment can have important consequences on the equilibrium properties [16]. The induced new conditions change the relative fitness of the species and, therefore, changes the final outcome of the selective process. This modification can stabilized the population by endowing the species with a larger survival probability or, on the contrary, it can destabilize the population by pushing the external temperature out of the limits of survival of the existing species. In the latter case, this induced-variation can drive the population to extinction. In any case, absolute stability does not exist in these models. The possibility of modifying the environment is an additional factor that contributes to keep open forever the fate of evolution.

